# PAM recognition by miniature CRISPR nucleases triggers programmable double-stranded DNA target cleavage

**DOI:** 10.1101/654897

**Authors:** Tautvydas Karvelis, Greta Bigelyte, Joshua K. Young, Zhenglin Hou, Rimante Zedaveinyte, Karolina Budre, Sushmitha Paulraj, Vesna Djukanovic, Stephen Gasior, Arunas Silanskas, Česlovas Venclovas, Virginijus Siksnys

## Abstract

In recent years, CRISPR-associated (Cas) nucleases have revolutionized the genome editing field. Being guided by an RNA to cleave double-stranded (ds) DNA targets near a short sequence termed a protospacer adjacent motif (PAM), Cas9 and Cas12 offer unprecedented flexibility, however, more compact versions would simplify delivery and extend application. Here, we present a collection of 10 exceptionally compact (422-603 amino acids) CRISPR-Cas nucleases that recognize and cleave dsDNA in PAM dependent manner. Categorized as class 2 type V-F they come from the Cas14 family and distantly related type V-U3 Cas proteins found in bacteria. Using biochemical methods, we demonstrate that a 5’ T- or C-rich PAM sequence triggers double stranded (ds) DNA target cleavage. Based on this discovery, we evaluated whether they can protect against invading dsDNA in *E. coli* and find that some but not all can. Altogether, our findings show that miniature Cas nucleases are functional CRISPR-Cas defense systems and have the potential to be harnessed as programmable nucleases for genome editing.

## INTRODUCTION

Clustered regularly interspaced short palindromic repeat (CRISPR) associated (Cas) microbial defense systems protect their hosts against foreign nucleic acid invasion (1–3). Utilizing small guide RNAs (gRNAs) transcribed from a CRISPR locus, accessory Cas proteins are directed to silence invading foreign RNA and DNA (2, 4, 5). Based on the number and composition of proteins involved in nucleic acid interference, CRISPR-Cas systems are categorized into distinct classes, 1-2, and types, I-VI (2, 6). Class 2 systems encode a single effector protein for interference and are further subdivided into types, II, V, and VI (7). Cas9 (type II) and Cas12 (type V) proteins have been shown to cleave invading double-stranded (ds) DNA, single-stranded (ss) DNA, and ssRNA (8–13). To recognize and cleave a dsDNA target, both Cas9 and Cas12 require a short sequence, termed the protospacer adjacent motif (PAM), in the vicinity of a DNA sequence targeted by the gRNA (8, 9, 14). Over the past several years, these endonucleases have been adopted as robust genome editing and transcriptome manipulation tools (15–18). Although both nucleases have been widely used, the size of Cas9 and Cas12 provides constraints on cellular delivery that may limit certain applications, including therapeutics (19, 20).

Recently, small CRISPR-associated effector proteins (Cas12f) belonging to the type V-F subtype have been identified through mining of sequence databases (6, 7, 13, 21). The majority of Cas12f proteins (including V-U3 and Cas14) are nearly half the size of the smallest Cas9 or Cas12 nucleases (Figure 1A) (2, 6, 7, 21). Compared to Cas12 orthologs, the N-terminal half of miniature Cas12f proteins differ significantly in length accounting for most of the size difference between the two groups (Fig. 1*A* and Supplementary Figure S1). Due to this and their similarity to transposase associated TnpB proteins, it is hypothesized that they are remnants or intermediates of type V CRISPR-Cas system evolution and are incapable of forming the protein architecture required for dsDNA target recognition and cleavage (7, 21). To our knowledge, functional characterization of Cas12f proteins are limited to Cas14a1 (Cas14 family) which were shown to exclusively target and cleave ssDNA in a PAM-independent manner (21). Additionally, target binding and cleavage by Cas14 triggers collateral nuclease activity that manifested as trans-acting non-specific ssDNA degradation (21). This seems to be a feature largely shared across the type V family, although Cas12g was reported to initially target ssRNA and then indiscriminately degrade both ssDNA and ssRNA (12, 13).

**Figure 1.**
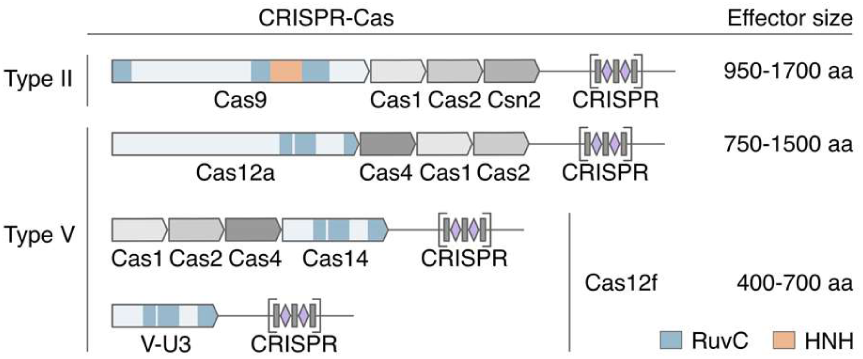
Schematic representation of CRISPR-Cas loci and effector proteins for Type II and V systems as exemplified by *Streptococcus pyogenes* SF370 (NC_002737.2) for Cas9, *Acidaminococcus sp*. BV3L6 (NZ_AWUR01000016.1) for Cas12a, uncultured archaeon (KU_516197.1) for Cas14 and *Syntrophomonas palmitatica* (NZ_BBCE01000017.1) for type V-U3, respectively. Tri-split RuvC domains of effector proteins are shown in blue, HNH domain of Cas9 in orange.

Here, using biochemical assays, we report that miniature Cas12f effectors (Cas14 and type V-U3), like most other Cas12 proteins, are able to cleave dsDNA targets if a 5’ PAM sequence is present in the vicinity of the guide RNA target. For the Cas14 family, this primarily consists of 5’ T-rich sequences while type V-U3 PAM recognition includes both C- and T-rich motifs. Additionally, we demonstrate that some type V-U3 bacterial nucleases (but none of the Cas14 family proteins tested) function as *bona fide* CRISPR-Cas systems in a *E. coli* cells to protect against invading dsDNA despite their small size. These findings provide the first experimental evidence that miniature type V-F effectors are programmable dsDNA nucleases and lay the foundation for the adoption of these proteins as genome editing tools.

## METHODS

### Engineering CRISPR-Cas12f systems encoding plasmids

Engineered CRISPR-Cas12f systems with three repeat:spacer:repeat units modified to target a 7 bp randomized PAM plasmid DNA library described previously (22) were synthesized (GenScript) and cloned into a modified pET-duet1 (MilliporeSigma) or pACYC184 (NEB) plasmids. For CRISPR-Cas14a1 system, pLBH531_MBP-Cas14a1 plasmid (gift from Jennifer Doudna, Addgene plasmid #112500) was used. The sequences of the Cas12f proteins are listed in Supplementary Data S1 file and the links to the plasmid sequences are provided in Supplementary Table S2.

### RNA synthesis

Cas14a1 single guide RNAs (sgRNA) were produced by *in vitro* transcription using TranscriptAid T7 High Yield Transcription Kit (Thermo Fisher Scientific) and purified using GeneJET RNA Purification Kit (Thermo Fisher Scientific). Templates for T7 transcription were generated by PCR using overlapping oligonucleotides, altogether, containing a T7 promoter at the proximal end followed by the sgRNA sequence. Sequences of the sgRNA used in our study are available in Supplementary Data S1 file.

### Detecting Cas12f dsDNA cleavage and PAM recognition

Plasmid DNA targets were cleaved with Cas12f ribonucleoprotein (RNP) complexes produced from the modified locus or by combining *E. coli* lysate containing Cas14a1 protein with T7 transcribed sgRNA. First*E. coli* DH5α or ArcticExpress (DE3) cells were transformed with CRISPR-Cas12f encoding plasmids (pACYC or pET-duet1 and pLBH531, respectively) and cultures grown in LB broth (30 ml) supplemented with either chloramphenicol (25 μg/ml) (pACYC plasmids) or ampicillin (100 μg/ml) (pET-duet1 and pLBH531 plasmids). Next, for plasmids with a T7 promoter (pET-duet1 and pLBH531 plasmids), expression was induced with 0.5 mM IPTG when cultures reached an OD_600_ of 0.5 and incubated overnight at 16°C. Cells (from 10 ml) were collected by centrifugation and re-suspended in 1 ml of lysis buffer (20 mM phosphate, pH 7.0, 0.5 M NaCl, 5% (v/v) glycerol) supplemented with 10 μl PMSF (final conc. 2 mM) and lysed by sonication. Cell debris was removed by centrifugation. 10 μl of the obtained supernatant containing RNPs were used directly in digestion experiments. For Cas14a1, 20 μl of clarified supernatant was combined with 1 μl of RiboLock RNase Inhibitor (Thermo Fisher Scientific) and 2 μg of sgRNA and allowed to complex with the clarified lysate as described below.

Cas12f RNP complexes were used to cleave either the 7 bp randomized PAM library or a plasmid containing a fixed PAM and gRNA target. Briefly, 10 μl of Cas12f-gRNA RNP containing lysate was mixed with 1 μg of PAM library or 1 μg of plasmid containing a single PAM and gRNA target in 100 μl of reaction buffer (10 mM Tris-HCl, pH 7.5 at 37°C, 100 mM NaCl, 1 mM DTT, and 10 mM MgCl2). After a 1 h incubation at 37°C, DNA ends were repaired by adding 1 μl of T4 DNA polymerase (Thermo Fisher Scientific) and 1 μl of 10 mM dNTP mix (Thermo Fisher Scientific) and incubating the reaction for 20 min at 11°C. The reaction was then inactivated by heating it up to 75°C for 10 min and 3’-dA overhangs added by incubating the reaction mixture with 1 μl of DreamTaq polymerase (Thermo Fisher Scientific) and 1 μl of 10 mM dATP (Thermo Fisher Scientific) for 30 min at 72°C. Additionally, RNA was removed by incubation for 15 min at 37°C with 1 μl RNase A (Thermo Fisher Scientific). Following purification with a GeneJet PCR Purification column (Thermo Fisher Scientific), the end repaired cleavage products (100 ng) were ligated with a double-stranded DNA adapter containing a 3’-dT overhang (100 ng) for 1 h at 22°C using T4 DNA ligase (Thermo Fisher Scientific). After ligation, cleavage products were PCR amplified appending sequences required for deep sequencing and subjected to Illumina sequencing (22, 23).

Double-stranded DNA target cleavage was evaluated by examining the unique junction generated by target cleavage and adapter ligation in deep sequence reads. This was accomplished by first generating a collection of sequences representing all possible outcomes of dsDNA cleavage and adapter ligation within the target region. For example, cleavage and adapter ligation at just after the 21st position of the target would produce the following sequence (5’-CCGCTCTTCCGATCTGCCGGCGACGTTGGGTCAACT-3’) where the adapter and target sequences comprise 5’-CCGCTCTTCCGATCT-3’ and 5’-GCCGGCGACGTTGGGTCAACT-3’, respectively. The frequency of the resulting sequences was then tabulated using a custom python script and compared to negative controls (experiments setup without functional Cas12f complexes) to identify target cleavage.

Evidence of PAM recognition was evaluated as described previously (22, 23). Briefly, the sequence of the protospacer adapter ligation exhibiting an elevated frequency in the previous step was used in combination with a 10 bp sequence 5’ of the 7 bp PAM region to identify reads that supported dsDNA cleavage. Once identified, the intervening 7 bp PAM sequence was isolated by trimming away the 5’ and 3’ flanking sequences using a custom python script and the frequency of the extracted PAM sequences normalized to the original PAM library to account for inherent biases using the following formula.

Normalized Frequency = (Treatment Frequency)/((Control Frequency)/(Average Control Frequency))

Following normalization, a position frequency matrix (PFM) (24) was calculated and compared to negative controls (experiments setup without functional Cas12f complexes) to look for biases in nucleotide composition as a function of PAM position. Biases were considered significant and indicative of PAM recognition if they deviated by more than 2.5-fold from the negative control. Analyses were limited to the top 10% most frequent PAMs to reduce the impact of background noise resulting from non-specific cleavage coming from other components in the *E. coli* cell lysate mixtures.

### Expression and purification of Cas14a1 proteins

Cas14a1 protein was expressed in *E. coli* BL21(DE3) strains from the pLBH531_MBP-Cas14a1 plasmid (gift from Jennifer Doudna, Addgene plasmid #112500). Cas14a1^D326A^ and Cas14a1^D510A^ expression plasmids were engineered from pLBH531 using Phusion Site-Directed Mutagenesis Kit (Thermo Fisher Scientific). *E. coli* cells were grown in LB broth supplemented with ampicillin (100 μg/ml) at 37°C. After culturing to an OD_600_ of 0.5, temperature was decreased to 16°C and expression induced with 0.5 mM IPTG for Cas14a1, Cas14a1^D326A^ and Cas14a1^D510A^. After 16 h cells were pelleted, re-suspended in loading buffer (20 Tris-HCl, pH 8.0 at 25°C, 1.5 M NaCl, 5 mM 2-mercaptoethanol, 10 mM imidazole, 2 mM PMSF, 5% (v/v) glycerol) and disrupted by sonication. Cell debris was removed by centrifugation. The supernatant was loaded on the Ni^2+^-charged HiTrap chelating HP column (GE Healthcare) and eluted with a linear gradient of increasing imidazole concentration (from 10 to 500 mM) in 20 Tris-HCl, pH 8.0 at 25°C, 0.5 M NaCl, 5 mM 2-mercaptoethanol buffer. The fractions containing Cas14a1 variants were pooled and subsequently loaded on HiTrap heparin HP column (GE Healthcare) for elution using a linear gradient of increasing NaCl concentration (from 0.1 to 1.5 M). The fractions containing protein were pooled and the 10×His-MBP-tag was cleaved by overnight incubation with TEV protease at 4°C. To remove cleaved the 10×His-MBP-tag and TEV protease recognition site, reaction mixtures were loaded onto a HiTrap heparin HP 5 column (GE Healthcare) for elution using a linear gradient of increasing NaCl concentration (from 0.1 to 1.5 M). Next, the elution from the HiTrap heparin column was loaded on a MBPTrap column (GE Healthcare) and Cas14a1 protein was collected in the flow-through. The collected fractions with Cas14a1 were then dialyzed against 20 mM Tris-HCl, pH 8.0 at 25°C, 500 mM NaCl, 2 mM DTT and 50% (v/v) glycerol and stored at −20°C. The sequences of the Cas14a1 proteins are listed in Supplementary Data S1 file.

### Cas14a1-sgRNA complex assembly for *in vitro* DNA cleavage

Cas14a1 ribonucleoprotein (RNP) complexes (1 μM) were assembled by mixing Cas14a1 protein with sgRNA at 1:1 molar ratio followed by incubation in a complex assembly buffer (10 mM Tris-HCl, pH 7.5 at 37°C, 100 mM NaCl, 1 mM EDTA, 1 mM DTT) at 37°C for 30 min.

### DNA substrate generation

Plasmid DNA substrates were generated by cloning oligoduplexes assembled after annealing complementary oligonucleotides (Metabion) into pUC18 plasmid over HindIII (Thermo Fisher Scientific) and EcoRI (Thermo Fisher Scientific) restriction sites. The sequences of the inserts are listed in Supplementary Data S1 file and the links to the plasmid sequences are provided in Supplementary Table S2.

To generate radiolabeled DNA substrates, the 5’-ends of oligonucleotides were radiolabeled using T4 PNK (Thermo Fisher Scientific) and [γ-33P]ATP (PerkinElmer). Duplexes were made by annealing two oligonucleotides with complementary sequences at 95°C following slow cooling to room temperature. A radioactive label was introduced at the 5’-end of individual DNA strands before annealing with the unlabeled strands. The sequences of the oligoduplexes are provided in Supplementary Data S1 file.

### DNA substrate cleavage assay

Plasmid DNA cleavage reactions were initiated by mixing plasmid DNA with Cas14a1 RNP complex at 46°C. The final reaction mixture typically contained 3 nM plasmid DNA, 100 nM Cas14a1 RNP complex in 2.5 mM Tris-HCl, pH 7.5 at 37°C, 25 mM NaCl, 0.25 mM DTT and 10 mM MgCl_2_ reaction buffer. Aliquots were removed at timed intervals (30 min if not indicated differently) and mixed with 3× loading dye solution (0.01% Bromophenol Blue and 75 mM EDTA in 50% (v/v) glycerol) and reaction products were analyzed by agarose gel electrophoresis and ethidium bromide staining.

Reactions with oligoduplexes were typically carried-out by mixing labelled oligoduplex with Cas14a1 RNP complex and incubating at 46°C. The final reaction mixture contained 1 nM labeled duplex, 100 nM Cas14a1 RNP complex, 5 mM Tris-HCl, pH 7.5 at 37°C, 50 mM NaCl, 0.5 mM DTT and 5 mM MgCl_2_ in a 100 μl reaction volume. Aliquots of 6 μl were removed from the reaction mixture at timed intervals (0, 1, 2, 5, 10, 15 and 30 min), quenched with 10 μl of loading dye (95% (v/v) formamide, 0.01% Bromophenol Blue and 25 mM EDTA) and subjected to denaturing gel electrophoresis (20% polyacrylamide containing 8.5 M urea in 0.5× TBE buffer). Gels were dried and visualized by phosphorimaging.

### M13 cleavage assay

M13 ssDNA cleavage reactions were initiated by mixing M13 ssDNA (New England Biolabs) and DNA activator with Cas14a1 RNP complex at 46°C. Cleavage assays were conducted in 2.5 mM Tris-HCl, pH 7.5 at 37°C, 25 mM NaCl, 0.25 mM DTT and 10 mM MgCl_2_. The final reaction mixture contained 5nM M13 ssDNA, 100 nM ssDNA or dsDNA activator and 100 nM Cas14a1 RNP. The reaction was initiated by addition of Cas14a1 RNP complex and was quenched at timed intervals (0, 5, 15, 30, 60 and 90 min) by mixing with 3× loading dye solution (0.01% Bromophenol Blue and 75 mM EDTA in 50% (v/v) glycerol). Products were separated on an agarose gel and stained with SYBR Gold (Thermo Fisher Scientific). The sequences of the activators are listed in Supplementary Data S1 file.

### Plasmid interference assay

Plasmid interference assays were performed in *E. coli* Arctic Express (DE3) strain bearing Cas12f systems (plasmids encoding CRISRP-Cas12f systems are listed in Supplementary Table S2). For Cas14a1, *E. coli* BL21 (DE3) strain was transformed with pGB53 plasmid, which was engineered from the pLBH545_Tet-Cas14a1_Locus plasmid (gift from Jennifer Doudna, Addgene plasmid #112501) by removing tracrRNA and CRISPR array with Bsp1407I and AvrII and adding sgRNA encoding sequence with T7 promoter, HDV ribozyme and terminator sequences. The cells were grown at 37°C to OD_600_ of ∼0.5 and prepared for chemical transformation with 100 ng of low copy number pSC101 target plasmids obtained by cloning oligoduplexes over EcoRI and XhoI or EcoRI and NheI restriction sites into pTHSSe_1 (gift from Christopher Voigt, Addgene plasmid #109233) or pSG4K5 (gift from Xiao Wang, Addgene plasmid #74492) plasmids, respectively (the sequences of the inserts are listed in Supplementary Data S1 file and the links to the plasmid sequences are provided in Supplementary Table S2). The co-transformed cells were further diluted by serial 10× fold dilutions and grown on plates containing inductor and antibiotics. For Cas14a1 – AHT (50 ng/ml), IPTG (0.5 mM) and chloramphenicol (30 μg/ml); Cas14b4 - gentamycin (10 μg/ml) and carbenicillin (100 μg/ml); for all other Cas12f proteins – IPTG (0.5 mM), gentamycin (10 μg/ml) and carbenicillin (100 μg/ml) at 37°C for 16–20 h.

## RESULTS

### DNA targeting requirements by Cas12f proteins

CRISPR-Cas14 is a recently identified class 2 type V-F CRISPR-Cas system (6, 21). In addition to encoding one of the smallest effector proteins described to date, these CRISPR systems also encode adaptation proteins (Cas1, Cas2 and Cas4) required for acquiring and incorporating new spacers into the CRISPR (Figure 1). To our knowledge, just a single Cas14 protein, Cas14a1, has been experimentally characterized where it was shown to only cleave ssDNA targets in a PAM-independent manner (21). In our experimentation, shared structural features with other type V effectors capable of dsDNA target cleavage (Supplementary Figure S1) motivated us to evaluate Cas14 family proteins for dsDNA cleavage activity. To establish DNA cleavage requirements, we initially tested the Cas14b4 protein (Supplementary Table S1) (21). First, spacers capable of targeting a randomized PAM plasmid library (22) were incorporated into its CRISPR array. The resulting modified CRISPR-Cas14b4 locus was synthesized, cloned into a low copy plasmid, and transformed into *E. coli*. A PAM determination assay (22, 23) was adapted to test the ability of the Cas14b4 to recognize and cleave a dsDNA target *in vitro* (Figure 2A). This was accomplished by combining clarified lysate containing Cas14b4 protein and gRNAs expressed from the reengineered locus with the PAM library. Next, DNA breaks were captured by double-stranded adapter ligation, enriched by PCR, and deep sequenced as described previously (22, 23). DNA cleavage occurring in the target sequence was evaluated by scanning regions in the protospacer for elevated frequencies of adapter ligation relative to negative controls (experiments using lysate from *E. coli* not transformed with the Cas14b4 locus). A slight increase in the number of adapter ligated sequences was recovered after the 21^st^ protospacer position 3’ of the randomized PAM (Supplementary Figure S2A). Analysis of these fragments showed the recovery of a T-rich sequence (5’-TTAT-3’) immediately 5’ of the gRNA target only in the Cas14b4 treated sample (Figure 2B and Supplementary Figure S2B). To confirm the PAM sequence and dsDNA cleavage position, a plasmid was constructed containing a target adjacent to the identified 5’-TTAT-3’ PAM sequence and subjected to cell lysate cleavage experiments. To increase Cas14b4 concentration in the cell lysate, a higher copy number DNA expression plasmid equipped with an inducible T7 promoter was also utilized. Sequencing of the target plasmid cleavage products confirmed cleavage at the 21^st^ position (Supplementary Figure S2C). Reactions using deletion variants further confirmed that Cas14b4 was the sole endonuclease required for the observed dsDNA target recognition and cleavage activity (Supplementary Figure S2C).

**Figure 2.**
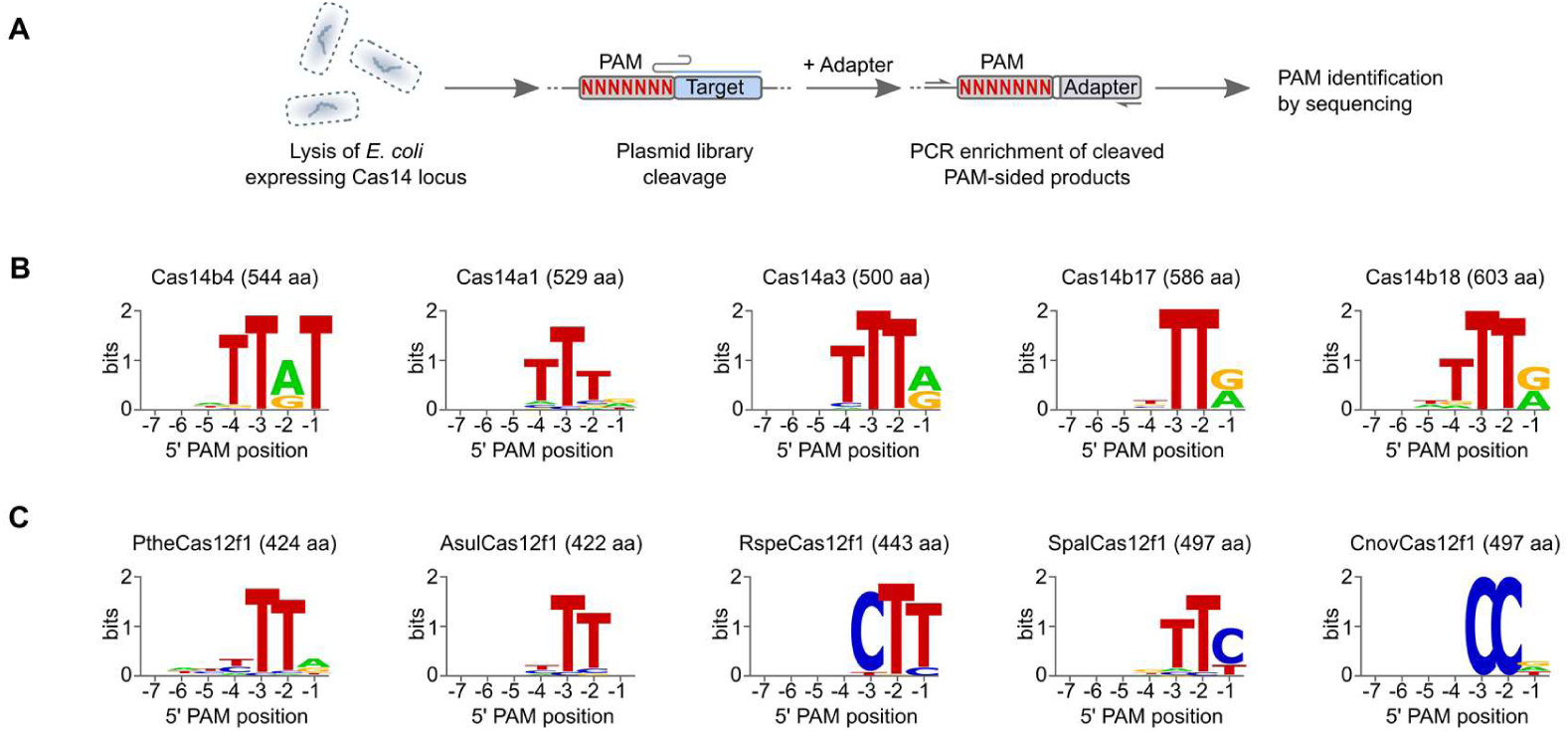
PAM sequences of Cas12f effector proteins. (**A**) Workflow of the biochemical approach used to examine PAM recognition and detect dsDNA cleavage. WebLogos of PAM sequences for Cas14 (**B**) and Cas12f1 (**C**) proteins.

Four additional Cas14 proteins were also evaluated. These consisted of Cas14a1, Cas14a3, Cas14b17, and Cas14b18 proteins (Supplementary Table S1). First, for Cas14a3, Cas14b17, and Cas14b18 a T7 inducible expression plasmids were synthesized. They contained a minimal locus: the *cas14* gene, sequence encoding a putative tracrRNA between the nuclease gene and CRISPR repeats, and a CRISPR array modified to target the PAM library. Then, similar to Cas14b4 experimentation, *E. coli* lysate from cells expressing the Cas14 nuclease and guide RNAs was combined with a randomized PAM library. Cleavage products were then captured and analyzed as described for Cas14b4 (Figure 2A). For Cas14a1, a modified approach was used. Here, an *in vitro* transcribed single guide RNA (sgRNA) capable of targeting the PAM library was combined with *E. coli* lysate containing Cas14a1 protein and then assayed for dsDNA target recognition and cleavage as described above. Similarly, to Cas14b4, a cleavage signal distal to the PAM region was detected for all proteins. For Cas14b nucleases, this occurred just after the 21^st^ protospacer position while for Cas14a proteins the signal was found after the 24^th^ position (Supplementary Figures S3A, S4A and S5A). Analysis of the sequences supporting cleavage yielded 5’ T-rich PAM recognition like that recovered for Cas14b4 (Figure 2C and Supplementary Figures S3B, S4B and S5B).

Next, to further explore the DNA cleavage requirements of miniature CRISPR-Cas effectors, we sought to evaluate the dsDNA cleavage activity of proteins from an uncharacterized putative group of class 2 CRISPR-Cas systems, type V-U3, which lack the proteins involved in adaptation (2, 7). First, PSI-BLAST searches were performed to identify a group of CRISPR-associated proteins primarily from bacteria, in particular, lineages of *Clostridia* and *Bacilli*, that belonged to the type V-U3 family (Supplementary Table S1). They contained a conserved C-terminal tri-split RuvC domain similar to Cas14 and other Cas12 nucleases and a short variable N-terminal sequence as observed for the Cas14 family. Next, five were selected for functional characterization. In general, they were chosen to represent the uncovered diversity and ranged in size between 422-497 amino acids. As described for Cas14 (Figure 2A), minimal CRISPR loci containing the nuclease gene, putative tracrRNA encoding region, and CRISPR array reprogrammed to target the PAM library were then synthesized and cloned into an inducible expression plasmid and examined for dsDNA target recognition and cleavage. As shown in Supplementary Figures S6-S10, all produced cleavage around the 24^th^ position 3’ of the PAM region. For nucleases from *Parageobacillus thermoglucosidasius* (Pthe) and *Acidibacillus sulfuroxidans* (Asul), secondary cleavage signals (>5% of all reads) were also recovered either before or after the 24^th^ position. Like Cas14, type V-U3 nucleases cleaved the dsDNA library in a 5’ PAM-dependent manner and altogether expanded miniature Cas nucleases PAM diversity to encompass not only T-rich but also C-rich motifs (Figure 2C).

### Biochemical characterization of Cas14a1 mediated dsDNA cleavage

Since a sgRNA solution was available for Cas14a1 (21) we further probed the Cas14a1 protein for programmable dsDNA cleavage using purified components. First, to decipher optimal reaction conditions, the effect of sgRNA spacer length, temperature, salt concentration, and divalent metal ions were evaluated on Cas14a1 dsDNA cleavage activity (Supplementary Figure S4). Experiments revealed that Cas14a1 ribonucleoprotein (RNP) complex is a Mg^2+^-dependent endonuclease that functions best in low salt concentrations (5-25 mM) and is active in a wide temperature range ∼37-50°C, with a temperature optimum of ∼46°C (Supplementary Figure S4). Furthermore, sgRNA spacers of around 20 nt supported the most robust dsDNA cleavage activity (Supplementary Figure S4). Under optimized reaction conditions, supercoiled (SC) plasmid DNA containing a target sequence flanked by a Cas14a1 PAM (5’-TTTA-3’) was completely converted to a linear form (FLL) indicating the formation of a dsDNA break (Figure 3A). Additionally, cleavage of linear DNA yielded DNA fragments of expected size, further validating Cas14a1 mediated dsDNA break formation. Next, we confirmed that Cas14a1 requires both PAM and sgRNA recognition to cleave a dsDNA target (Supplementary Figure S7). Finally, alanine substitution of conserved RuvC active site residues abolished cutting activity, confirming that the RuvC domain is essential for the observed dsDNA cleavage activity (Figure 3B).

**Figure 3.**
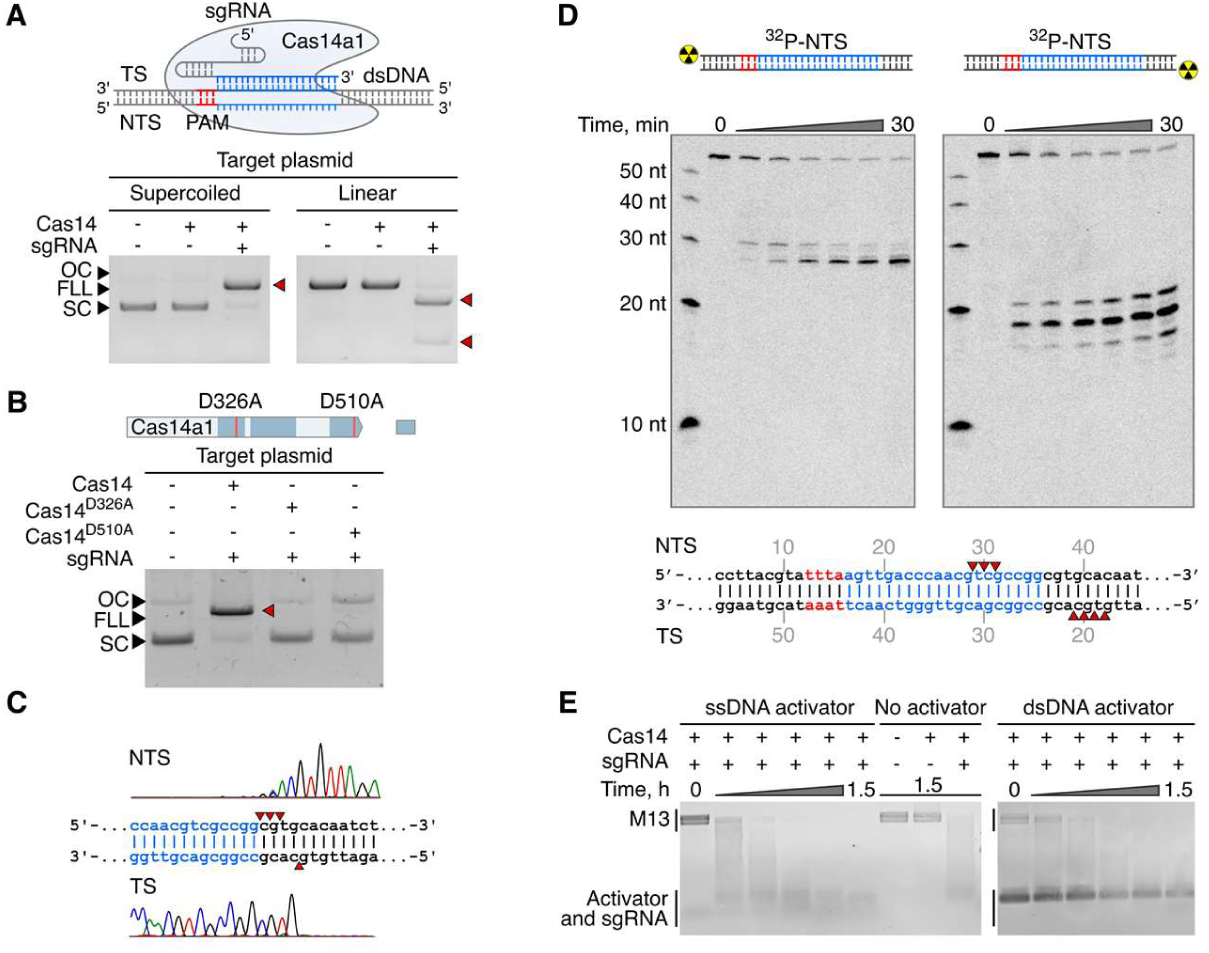
Cas14a1 RNP complex is a PAM-dependent dsDNA endonuclease. (**A**) Cas14a1 RNP complex cleaves plasmid DNA targets *in vitro* in a PAM-dependent manner. (**B**) Alanine substitution of two conserved RuvC active site residues completely abolishes Cas14a1 DNA cleavage activity. (**C**) Run-off sequencing of Cas14a1 pre-cleaved plasmid DNA indicates that cleavage is centered around positions 20-24 bp 3’ of the PAM resulting in 5’ overhangs. (**D**) Oligoduplex cleavage patterns are consistent with staggered cleavage but differ from experiments assembled with plasmid DNA (shown in C). (**E**) Collateral non-specific M13 ssDNA degradation activity by Cas14a1 triggered by ssDNA and PAM-containing dsDNA. TS – target strand, NTS – non-target strand, SC – supercoiled, FLL – full length linear, OC – open circular.

The type of dsDNA break generated by Cas14a1 was examined next. Using run-off sequencing, we observed that Cas14a1 makes 5’ staggered overhanging DNA cut-sites. Cleavage predominantly occurred centered around positions 20-24 bp in respect to PAM sequence (Figure 3C) and was independent of spacer length or plasmid topology (Supplementary Figure S8). The cleavage pattern of Cas14a1 was also assessed on synthetic double-stranded oligodeoxynucleotides. As illustrated in Figure 3D and Supplementary Figure S9, a 5’staggered cut pattern, albeit with less strictly defined cleavage positions than observed with larger DNA fragments was seen.

Next we investigated if the non-specific ssDNA degradation activity of Cas14a1 could be induced not just by ssDNA targets (21) but also by dsDNA targets. First, the ability of Cas14a1 to indiscriminately degrade single-stranded M13 DNA in the presence of a ssDNA target without a PAM was confirmed. Then, a dsDNA target containing a 5’ PAM and sgRNA target for Cas14a1 was also tested for its ability to trigger non-selective ssDNA degradation. As shown in Figure 3E, the trans-acting ssDNase activity of Cas14a1 was activated by both ssDNA and dsDNA targets, similar to observations made for Cas12a (12).

### Cas12f mediated plasmid DNA interference in *E. coli*

We next tested if Cas12f (Cas14 and type V-U3) systems can be programmed to target and cleave invading dsDNA in a heterologous *E. coli* host. First, an *E. coli* plasmid DNA interference assay was adopted (25, 26) using a minimal Cas14 or type V-U3 CRISPR locus modified to target the incoming low copy number plasmid DNA. To assess transformation efficiency, each experiment was serially diluted by 10× and compared with controls (experiments performed with a plasmid that does not contain a target site). Surprisingly, none of the Cas14 nucleases interfered with plasmid DNA transformation as evidenced by similar recovery of resistant colonies compared to controls (Figure 4A). These findings, however, are in agreement with a previous study which showed that Cas14a1 was incapable of depleting PAM plasmid libraries in a heterologous *E. coli* host and presumably due to this reason failed to detect PAM-dependent dsDNA cleavage (21). In contrast, type V-U3 effectors from *A. sulfuroxidans* (Asul) and *Syntrophomonas palmitatica* (Spal) both produced notable levels of plasmid interference and slight interference was also observable for nucleases from *P. thermoglucosidasius* (Pthe) and *Ruminococcus* sp. (Rspe) (Figure 4B).

**Figure 4.**
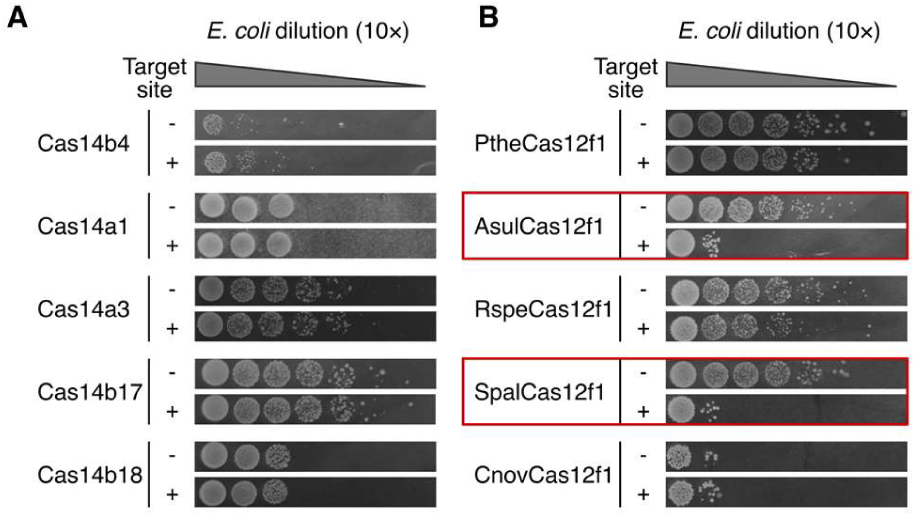
Cas12f mediated plasmid DNA interference in *E. coli*. (**A**) Plasmid interference assay for Cas14 and type V-U3 variants (**B**). Cells bearing a minimal CRISPR-Cas12f locus were transformed with a low copy number plasmid DNA containing the target sequence. To assess transformation efficiency, each experiment was serially diluted by 10× and compared with controls (experiments performed with a plasmid that does not contain a target site). Red boxes indicate Cas12f1 variants that showed visually detectable levels of DNA interference activity.

## DISCUSSION

Contrary to previous hypotheses suggesting that Cas14 and type V-U3 CRISPR systems lack the ability to defend against invading dsDNA (7, 21), we illustrate that despite their miniature size some of these nucleases have this capacity. Here, using biochemical approaches, we provide evidence demonstrating that these compact proteins are programmable enzymes capable of introducing targeted dsDNA breaks like larger effectors, Cas9 and Cas12. First, we uncovered PAM-dependent dsDNA cleavage activity similar to other type V interference proteins (13, 14, 27) for 10 Cas12f family members. Then, we purified the nuclease and sgRNA for Cas14a1 and reconstituted its nuclease activity *in vitro* confirming that both PAM and sgRNA recognition are required for dsDNA target cleavage. Additionally, we observed that ssDNA collateral nuclease activity was triggered not only by ssDNA targets (21) but also by dsDNA targets, a feature shared by most other type V family members (12, 13). Finally, using plasmid DNA interference assays, we report that like shown previously (21), Cas14 failed to protect against invading dsDNA in *E. coli*, while even more compact CRISPR-associated type V-U3 nucleases, Asul (422 aa) and Spal (497 aa), efficiently interfered with plasmid transformation. Taken together, this confirms that at least some Cas12f effectors function like Cas9 and Cas12 nucleases to recognize, cleave, and protect against dsDNA invaders in a heterologous host. In all, our results significantly improve our understanding of novel CRISPR-Cas systems and pave the way for the adoption of programmable miniature nucleases for genome editing applications.

## ACCESSION NUMBERS

Sequencing data are available on the NCBI Sequence Read Archive under BioProject ID <u>PRJNA607069</u>.

## SUPPLEMENTARY DATA

Supplementary Data are available on bioRxiv online.

## ACKNOWLEDGEMENT

We would like to thank S. Dooley, H. Lin, and M. King for informatics support and C. Skrdlant, and G. Zastrow-Hayes for Illumina sequencing support.

