## Supplemental Information for "PAM recognition by miniature CRISPR nucleases triggers programmable double-stranded DNA target cleavage"

**This PDF file includes:**

Supplementary Tables S1-S2

Supplementary Figures S1-S15

Legend Supplementary Data S1

SI References

**Other supplementary materials for this manuscript include the following:**

Supplementary Data S1

**Supplementary Table S1**. Cas14 and Cas12f1 proteins used in this study.

| Name | Size (aa) | Molecular mass (kDa) | Protein id (NCBI) | Organism | Scaffold accession (NCBI) |
|---|---|---|---|---|---|
| Cas14b4 | 544 | 63.9 | OIO21000.1 | Candidatus Micrarchaeota archaeon | MK005740.1 |
| Cas14a1 | 529 | 61.5 | QBM01166.1 | uncultured archaeon | MK005734 |
| Cas14a3 | 500 | 58.4 | QBM01093.1 | uncultured archaeon | MK005732 |
| Cas14b17* | 586 | 69.4 | RLG21245.1 | Candidatus Micrarchaeota archaeon | QMVG01000004.1 |
| Cas14b18* | 603 | 69.9 | RJP56748.1 | *Candidatus Aureabacteria bacterium* | QZJZ01000091.1 |
| PtheCas12f1[+] | 424 | 49.5 | WP_064552366.1 | *Parageobacillus thermoglucosidasius* | NZ_LXMA01000038.1 |
| AsulCas12f1[+] | 422 | 48.7 | WP_109431741.1 | *Acidibacillus sulfuroxidans* | NZ_MPDK01000047.1 |
| RspeCas12f1[+] | 440 | 51.2 | WP_117896622.1 | *Unclassified Ruminococcus* | NZ_QTWX01000005.1 |
| SpalCas12f1[+] | 497 | 56.9 | WP_054696859.1 | *Syntrophomonas palmitatica* | NZ_BBCE01000017.1 |
| CnovCas12f1[+] | 497 | 58.5 | WP_120361969.1 | *Clostridium novyi* | NZ_CP029458.1 |

∗ Identified by BLAST alignments against the NCBI NR database. Numbering is continued from that described in Harrington *et al*. 2018 (1).

[+] Type V-U3 nuclease identified by PSI-BLAST.

**Supplementary Table S2**. Plasmid map links.

| Plasmid name | Description | Link |
|---|---|---|
| PV424 | Cas14b4 engineered and intact locus (native expression) | https://benchling.com/s/seq-uDaLUdexDYQQSz7QDMXF |
| PV477 | Disrupted Cas14b4 (native expression) | https://benchling.com/s/seq-y7Fh5ryHUNNX7R3sDYMa |
| R-652 | Cas14b4 intact locus (T7 expression) | https://benchling.com/s/seq-DNGdXS8DCheZt5Nv6kMu |
| R-656 | Cas14b4 minimal locus (T7 expression) | https://benchling.com/s/seq-kUL4zwpBZDDJIwIwwLBT |
| R-658 | Minus Cas14b4 (T7 expression) | https://benchling.com/s/seq-A2E4WYBe0vLNbHiWQ2py |
| pLBH531 | 10×His-MBP-Cas14a1 expression | https://www.addgene.org/112500/ |
| pLBH545 | Cas14a1 locus (tetracycline inducible expression) | https://www.addgene.org/112501/ |
| pGB53 | Cas14a1 and sgRNA expression (pLBH545-based; T7 and tetracycline inducible expression) | https://benchling.com/s/seq-QoHpAbpI97JLSnhzh8Mn |
| pGB49 | 10×His-MBP-Cas14a1 D326A expression (pLBH531-based) | https://benchling.com/s/seq-qpUDPxM6IBhTJFXpNVUJ |
| pGB50 | 10×His-MBP-Cas14a1 D510A expression (pLBH531-based) | https://benchling.com/s/seq-W1miOvRPGZ44fgZnV0Nn |
| pCas14a3-pETduet-1 | Cas14a3 intact locus (T7 expression) | https://benchling.com/s/seq-6PdzgqR721Th21GAzyM7 |
| pCas14b17-pETduet-1 | Cas14b17 intact locus (T7 expression) | https://benchling.com/s/seq-QJzxQ4p6OonoQ8ZszpF7 |
| pCas14b18-pETduet-1 | Cas14b18 intact locus (T7 expression) | https://benchling.com/s/seq-6lJN8dk7ZTmbimkqw7Dg |
| pPtheCas12f1-pETduet-1 | PtheCas12f1 intact locus (T7 expression) | https://benchling.com/s/seq-qaU17VDPHblLUii1K5QZ |
| pAsulCas12f1-pETduet-1 | AsulCas12f1 intact locus (T7 expression) | https://benchling.com/s/seq-JWWCYPn66yxJI12WMBMX |
| pRspeCas12f1-pETduet-1 | RspeCas12f1 intact locus (T7 expression) | https://benchling.com/s/seq-XFA0y65xFCT57R719yVV |
| pSpalCas12f1-pETduet-1 | SpalCas12f1 intact locus (T7 expression) | https://benchling.com/s/seq-7LSUEFWIvEQk2AMpvsGA |
| pCnovCas12f1-pETduet-1 | CnovCas12f1 intact locus (T7 expression) | https://benchling.com/s/seq-92ITTuoZVYrJfN1hYgX1 |
| pTZ57 | 7N PAM plasmid library | https://benchling.com/s/seq-nu2IvfXbn7smVQ7T6MYi |
| pGB33 | Cas14b4 target plasmid (pUC18-based) | https://benchling.com/s/seq-iYcV6jflHOPbUxMdCGs9 |
| pGB40 | Cas14a1 target plasmid (pUC18-based) | https://benchling.com/s/seq-XGplLg5diY1G7BwBDHXn |

| | | |
|---|---|---|
| pGB41 | Cas14a1 ΔPAM (target plasmid) (pUC18-based) | https://benchling.com/s/seq-QS7mA2qo4e3JRlBvKsXb |
| pGB42 | Cas14a1 non-target plasmid (pUC18-based) | https://benchling.com/s/seq-rXjk6jlGlmPUSbb2GeT9 |
| pTHSSe_1 | Cas14a1 and Cas14b4 non-target plasmid (pSC101 ori) | https://www.addgene.org/109233/ |
| pGB43 | Cas14a1 target plasmid (pTHSSe_1-based) | https://benchling.com/s/seq-QJzxQ4p6OonoQ8ZszpF7 |
| pKP17 | Cas14b4 target plasmid (pTHSSe_1-based) | https://benchling.com/s/seq-g4SqeTyluv6C5FEwfmfS |
| pSG4K5 | Cas14a3, Cas14b17, Cas14b18 and all Cas12f1 non-target plasmid (pSC101 ori) | https://www.addgene.org/74492/ |
| pKP8 | Cas14a3, Cas14b17, Cas14b18, PtheCas12f1 and AsulCas12f1 target plasmid (pSG4K5-based) | https://benchling.com/s/seq-P3Bzoe526M7vh0Cv3mtD |
| pKP9 | RspeCas12f1 target plasmid (pSG4K5-based) | https://benchling.com/s/seq-8JkUE7ndI46oxuapcPWS |
| pKP10 | SpalCas12f1 target plasmid (pSG4K5-based) | https://benchling.com/s/seq-dIsHC0DVhRF4z8bLWdbw |
| pKP11 | CnovCas12f1 target plasmid (pSG4K5-based) | https://benchling.com/s/seq-6t7yhm7Vs6FwSj2N2qOu |

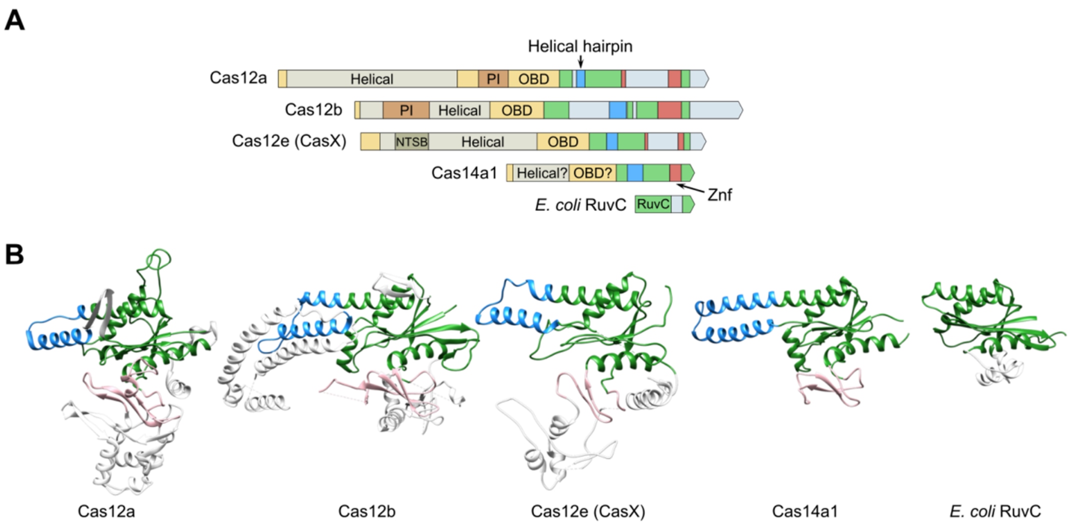


**Supplementary Figure S1**. Protein architecture and C-terminal RuvC domain structural comparisons between Cas12 dsDNA effectors and Cas14. Comparison includes Cas12a (PDB id: 5xut), Cas12b (PDB id: 5wti), Cas12e (CasX) (PDB id: 6ny2), Cas14a1 (model), and *E. coli* RuvC (PDB id: 1hjr). Cas14a1 region homologous to RuvC was identified using HHpred (2). Structural model for Cas14a1 sequence region was generated by the Rosetta comparative modeling protocol (3) using the structure of Cas12b (PDB id: 5WTI) as a modeling template. Five models generated by Rosetta were evaluated using the VoroMQA web server (4), designed to assess three-dimensional structures of proteins and protein complexes. The best-scoring model was selected and subjected to the protein structure refinement using the GalaxyRefine2 web server with standard refinement parameters (5). Multiple refined structural models returned by GalaxyRefine2 were again scored with VoroMQA and the best-scoring one was selected as the final model. (**A**) Organization of protein domains and relative size comparison between known type V dsDNA effectors and Cas14. (**B**) C-terminal RuvC domain of known type V dsDNA effectors, Cas14, and *E. coli* RuvC. The common RuvC core is colored green in all the structures. Helical hairpin common to Cas proteins, but absent from the *E. coli* RuvC is shown in blue. Common zinc finger motif (CasX and Cas14a1) or a zinc finger-like motif (Cas12a and Cas12b) is shown in pink. Additional structure-specific motifs are shown in grey.

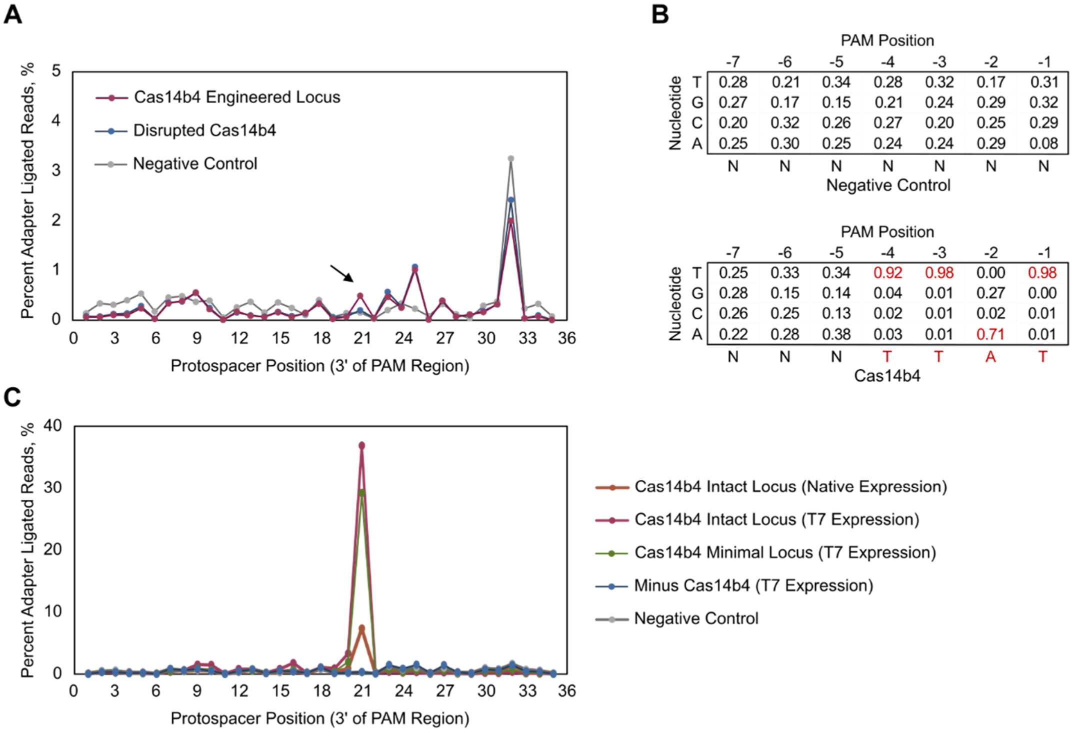


Negative Control (PAM Position)

| Nucleotide | -7 | -6 | -5 | -4 | -3 | -2 | -1 |
|---|---|---|---|---|---|---|---|
| T | 0.28 | 0.21 | 0.34 | 0.28 | 0.32 | 0.17 | 0.31 |
| G | 0.27 | 0.17 | 0.15 | 0.21 | 0.24 | 0.29 | 0.32 |
| C | 0.20 | 0.32 | 0.26 | 0.27 | 0.20 | 0.25 | 0.29 |
| A | 0.25 | 0.30 | 0.25 | 0.24 | 0.24 | 0.29 | 0.08 |
| | N | N | N | N | N | N | N |

Cas14b4 (PAM Position)

| Nucleotide | -7 | -6 | -5 | -4 | -3 | -2 | -1 |
|---|---|---|---|---|---|---|---|
| T | 0.25 | 0.33 | 0.34 | 0.92 | 0.98 | 0.00 | 0.98 |
| G | 0.28 | 0.15 | 0.14 | 0.04 | 0.01 | 0.27 | 0.00 |
| C | 0.26 | 0.25 | 0.13 | 0.02 | 0.01 | 0.02 | 0.01 |
| A | 0.22 | 0.28 | 0.38 | 0.03 | 0.01 | 0.71 | 0.01 |
| | N | N | N | T | T | A | T |

**Supplementary Figure S2**. Cas14b4 dsDNA recognition and cleavage. (**A**) Relative to the negative controls, the Cas14b4 locus engineered to target a PAM library produced a spike in the recovery of protospacer fragments ligated to an adapter just after the 21$^{st}$ position 3' of the PAM region. (**B**) PAM sequences that supported cleavage generated a position frequency matrix (PFM) exhibiting preferences for T and A bps 5' of the gRNA target. As a reference, a PFM at the same position in the lysate only control was also calculated. (**C**) dsDNA plasmids containing a PAM and gRNA target showed an even greater enrichment in the recovery of adapters ligated just after the 21$^{st}$ position, especially, for reactions where expression was enhanced with a T7 promoter. Experiments deleting *cas1*, *cas2*, and *cas4* genes (Cas14b4 Minimal Locus) and the *cas14b4* gene itself (Minus Cas14b4) confirmed that Cas14b4 was the only protein required for the observed dsDNA target recognition and cleavage.

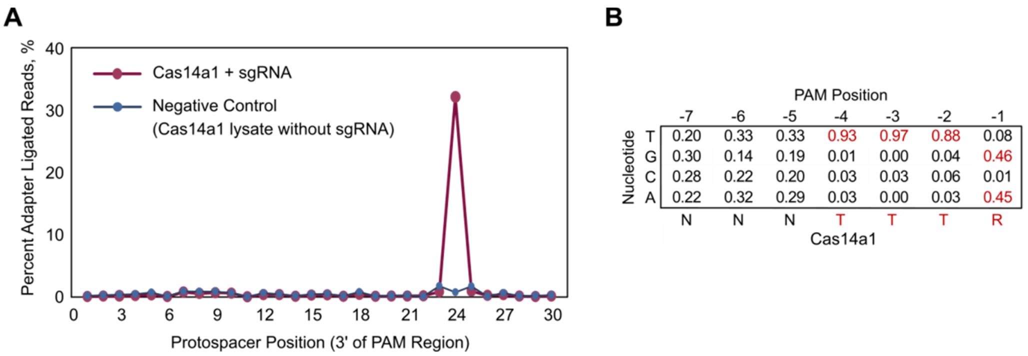


**Supplementary Figure S3**. Cas14a1 dsDNA recognition and cleavage. (**A**) *E. coli* lysate containing Cas14a1 and sgRNA targeting the PAM library produced an enrichment in the recovery of protospacer adapter ligated fragments just after the 24th position 3’ of the PAM region relative to the negative control. (**B**) Displayed as a PFM, PAM sequences that supported cleavage showed a 5’ T-rich PAM.

**A**

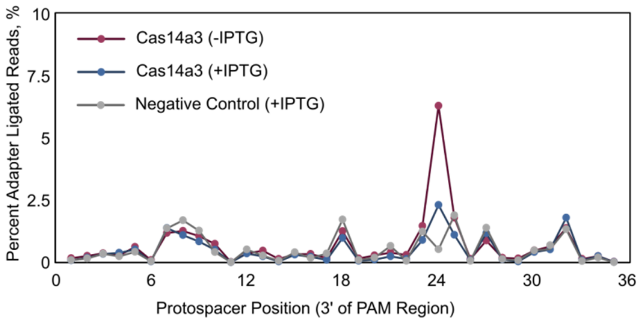


**B**

PAM Position

| Nucleotide | -7 | -6 | -5 | -4 | -3 | -2 | -1 |
|---|---|---|---|---|---|---|---|
| T | 0.21 | 0.24 | 0.34 | 0.88 | 1.00 | 0.98 | 0.00 |
| G | 0.28 | 0.22 | 0.15 | 0.02 | 0.00 | 0.00 | 0.42 |
| C | 0.26 | 0.24 | 0.20 | 0.07 | 0.00 | 0.01 | 0.02 |
| A | 0.26 | 0.30 | 0.31 | 0.03 | 0.00 | 0.01 | 0.56 |
| Cas14a3 | N | N | N | T | T | T | R |

**Supplementary Figure S4**. Cas14a3 dsDNA recognition and cleavage. (**A**) *E. coli* lysate from cells expressing the minimal CRISPR-Cas14a3 locus modified to target the 7N PAM library produced a spike in the recovery of adapter ligated fragments just after the 24$^{th}$ position 3' of the PAM with and without induction of expression with IPTG. (**B**) Similar to Cas14a1, a 5' T-rich PAM was shown to coincide with the cleavage signal.

A

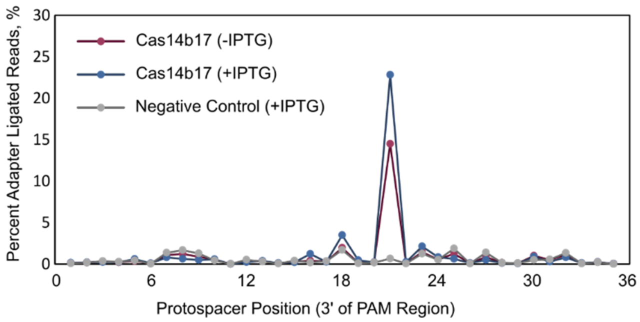


B

PAM Position

| Nucleotide | -7 | -6 | -5 | -4 | -3 | -2 | -1 |
|---|---|---|---|---|---|---|---|
| T | 0.23 | 0.27 | 0.30 | 0.47 | 0.99 | 0.99 | 0.01 |
| G | 0.26 | 0.18 | 0.23 | 0.21 | 0.00 | 0.00 | 0.53 |
| C | 0.27 | 0.27 | 0.25 | 0.19 | 0.01 | 0.01 | 0.02 |
| A | 0.24 | 0.27 | 0.22 | 0.13 | 0.00 | 0.00 | 0.45 |
| Cas14b17 | N | N | N | N | T | T | R |

**Supplementary Figure S5**. Cas14b17 dsDNA recognition and cleavage. (**A**) *E. coli* lysate containing Cas14b17 protein and guide RNAs targeting the PAM library produced an enrichment in the recovery of protospacer adapter ligated fragments just after the 21$^{st}$ position 3’ of the PAM region relative to the negative control (with and without IPTG induction of expression). (**B**) Displayed as a PFM, fragments with an adapter ligated after the 21$^{st}$ position exhibited a strong bias towards 5’-TTR(A/G)-3’ sequences in the PAM library.

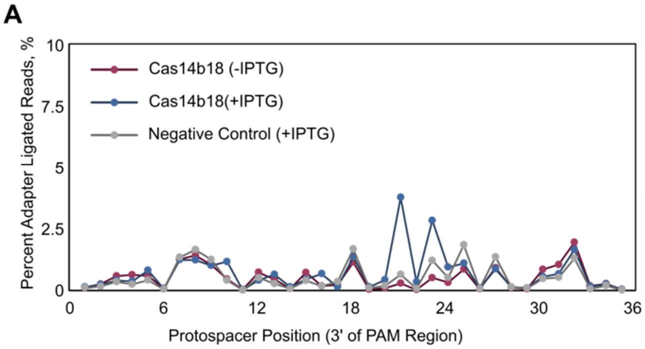


**B**

PAM Position

| Nucleotide | -7 | -6 | -5 | -4 | -3 | -2 | -1 |
|---|---|---|---|---|---|---|---|
| T | 0.25 | 0.29 | 0.42 | 0.81 | 1.00 | 0.97 | 0.02 |
| G | 0.22 | 0.15 | 0.11 | 0.08 | 0.00 | 0.00 | 0.55 |
| C | 0.24 | 0.22 | 0.13 | 0.03 | 0.00 | 0.02 | 0.00 |
| A | 0.29 | 0.35 | 0.35 | 0.08 | 0.00 | 0.00 | 0.43 |
| | N | N | N | T | T | T | R |

Cas14b18

**Supplementary Figure S6**. Cas14b18 dsDNA recognition and cleavage. (**A**) *E. coli* lysate containing Cas14b18 and guide RNAs targeting the PAM library produced an enrichment in the recovery of protospacer adapter ligated fragments just after the 21st position 3’ of the PAM region relative to the negative control. (**B**) Displayed as a PFM, fragments with an adapter ligated after the 21st position exhibited a strong bias towards 5’-TTTR(A/G)-3’ sequences in the PAM library.

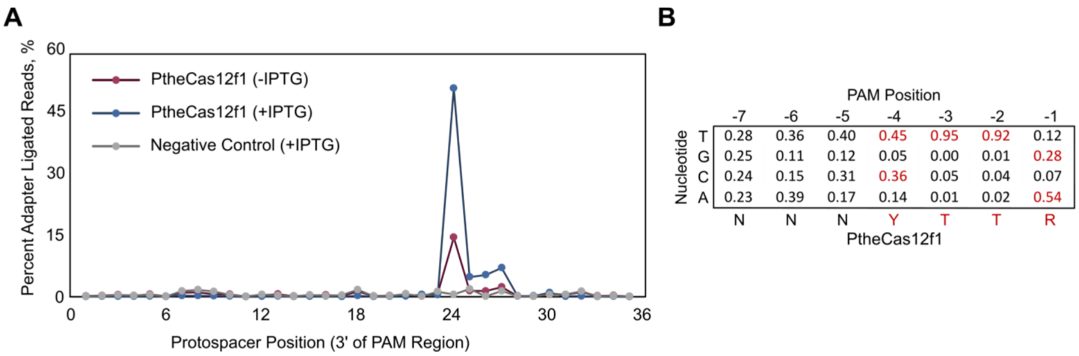


**Supplementary Figure S7**. PtheCas12f1 dsDNA recognition and cleavage. (**A**) *E. coli* lysate from cells expressing the minimal CRISPR-PtheCas12f1 locus modified to target the 7N PAM library produced a spike in the recovery of adapter ligated fragments just after the 24$^{th}$ position 3' of the PAM with and without induction of expression with IPTG. (**B**) Displayed as a PFM, fragments with an adapter ligated after the 24$^{th}$ position showed preference for 5'-Y(T/C)TTR(A/G)-3' sequences in the PAM library.

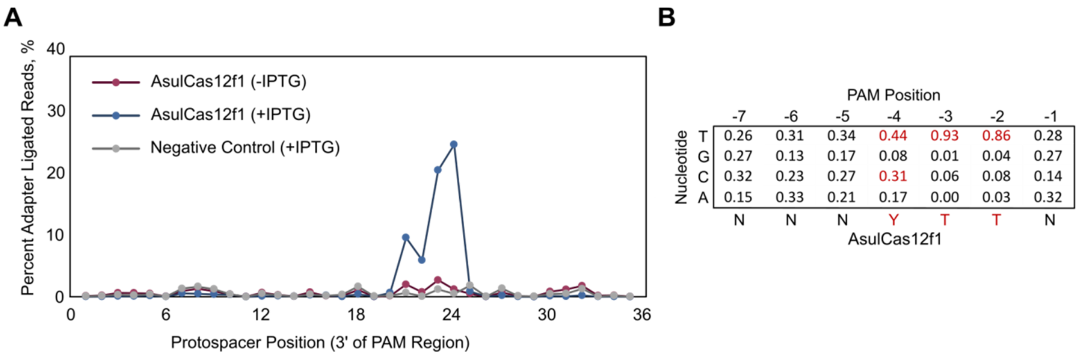


**Supplementary Figure S8**. AsulCas12f1 dsDNA recognition and cleavage. (**A**) Relative to the negative control, the AsulCas12f1 locus engineered to target a PAM library produced a spike in the recovery of protospacer fragments ligated to an adapter after the 21$^{st}$ position 3' of the PAM region. (**B**) Displayed as a PFM, fragments with an adapter ligated after the 21$^{st}$ position revealed a strong bias towards 5'-Y(T/C)TT-3' sequences in the PAM library.

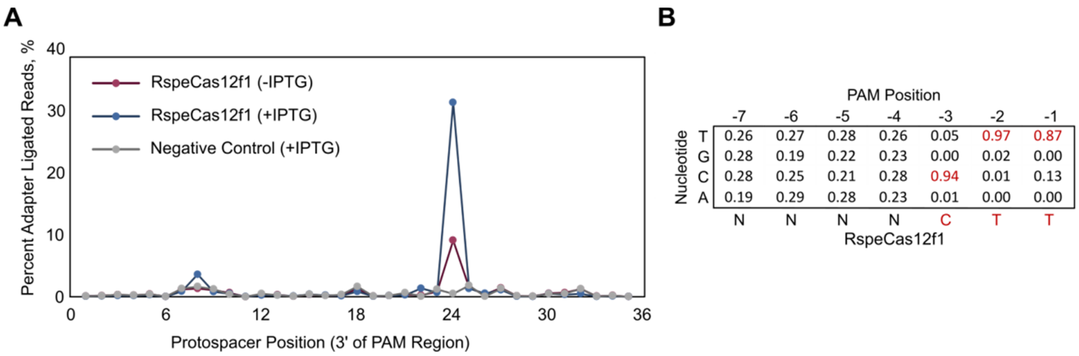


**Supplementary Figure S9**. RspeCas12f1 dsDNA recognition and cleavage. (**A**) *E. coli* lysate containing RspeCas12f1 protein and guide RNAs targeting the PAM library produced an enrichment in the recovery of protospacer adapter ligated fragments just after the 24$^{th}$ position 3' of the PAM region relative to the negative control (with and without IPTG induction of expression). (**B**) Displayed as a PFM, fragments with an adapter ligated after the 24$^{th}$ position demonstrated preference for 5'-CTT-3' sequences in the PAM library.

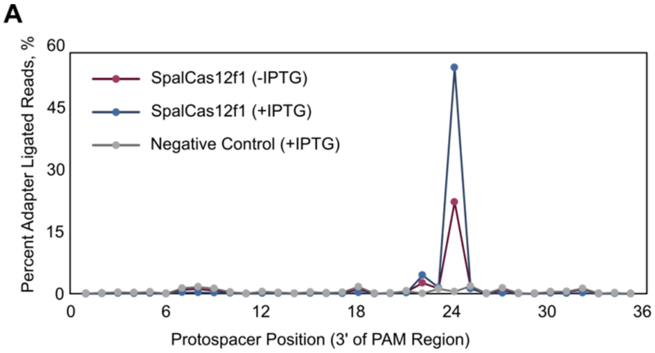


**B**

PAM Position

| Nucleotide | -7 | -6 | -5 | -4 | -3 | -2 | -1 |
|---|---|---|---|---|---|---|---|
| T | 0.26 | 0.28 | 0.20 | 0.24 | 0.84 | 0.93 | 0.21 |
| G | 0.28 | 0.16 | 0.28 | 0.44 | 0.03 | 0.01 | 0.02 |
| C | 0.29 | 0.22 | 0.29 | 0.12 | 0.06 | 0.05 | 0.74 |
| A | 0.17 | 0.34 | 0.22 | 0.21 | 0.07 | 0.01 | 0.03 |
| SpalCas12f1 | N | N | N | N | T | T | C |

**Supplementary Figure S10**. SpalCas12f1 dsDNA recognition and cleavage. (**A**) *E. coli* lysate from cells expressing the minimal CRISPR-SpalCas12f1 locus modified to target the 7N PAM library produced a spike in the recovery of adapter ligated fragments just after the 24th position 3’ of the PAM with and without induction of expression with IPTG. (**B**) Displayed as a PFM, fragments with an adapter ligated after the 24th position exhibited a strong bias towards 5’-TTC-3’ sequences in the PAM library.

**A**

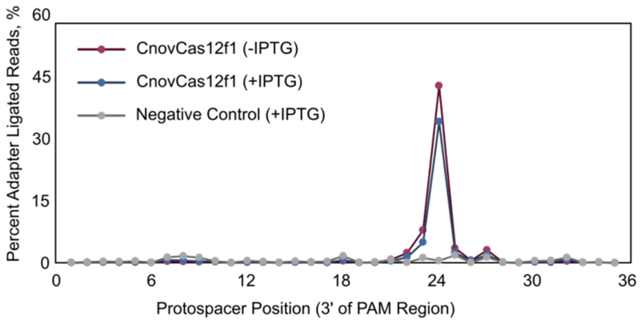


**B**

PAM Position

| Nucleotide | -7 | -6 | -5 | -4 | -3 | -2 | -1 |
|---|---|---|---|---|---|---|---|
| T | 0.27 | 0.26 | 0.25 | 0.25 | 0.00 | 0.00 | 0.30 |
| G | 0.25 | 0.25 | 0.24 | 0.23 | 0.00 | 0.00 | 0.35 |
| C | 0.25 | 0.24 | 0.27 | 0.26 | 1.00 | 1.00 | 0.02 |
| A | 0.23 | 0.25 | 0.23 | 0.25 | 0.00 | 0.00 | 0.33 |
| CnovCas12f1 | N | N | N | N | C | C | D |

**Supplementary Figure S11**. CnovCas12f1 dsDNA recognition and cleavage. (**A**) Relative to the negative controls, the CnovCas12f1 locus engineered to target a PAM library produced a spike in the recovery of protospacer fragments ligated to an adapter after the 24$^{th}$ position 3' of the PAM region. (**B**) Displayed as a PFM, fragments with an adapter ligated after the 24$^{th}$ position exhibited a strong bias towards 5'-CCD(T/G/A)-3' sequences in the PAM library.

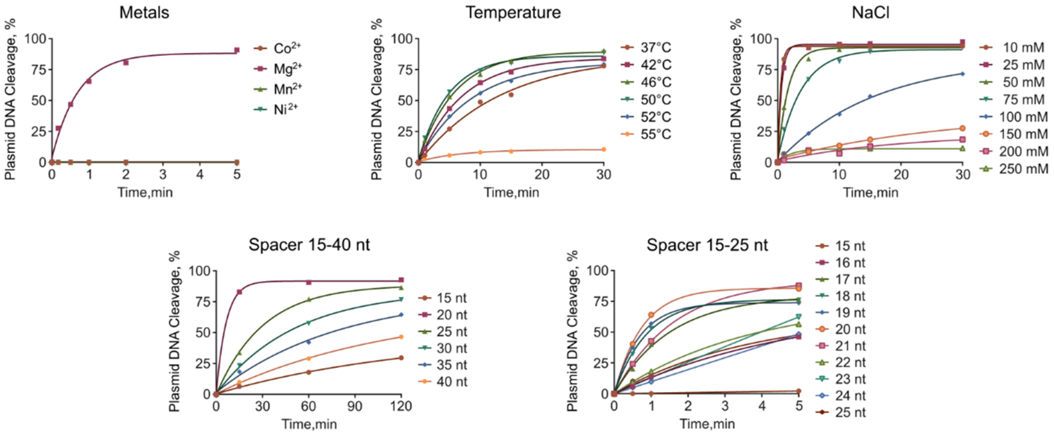


**Supplementary Figure S12**. Optimization of reaction conditions for Cas14a1 RNP mediated plasmid DNA cleavage. Cas14a1 RNP plasmid DNA cleavage was assayed by independently varying divalent metal ions, temperature, NaCl concentration, and sgRNA spacer length.

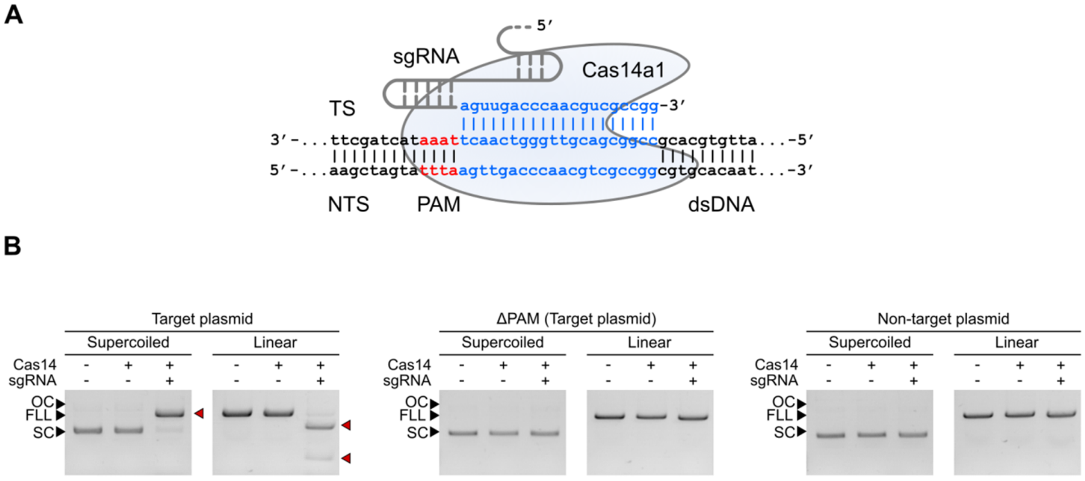


**Supplementary Figure S13**. Cas14a1 RNP complex is a PAM-dependent dsDNA endonuclease. (**A**) Schematic representation of Cas14a1 RNP complex with targeted DNA. (**B**) Cas14a1 RNP complexes cleaved plasmid DNA targets in a PAM-dependent manner (left panel) requiring both PAM (center panel) and sgRNA recognition (right panel). SC, OC, and FLL stand for supercoiled, open-circle, and full-length linearization, respectively. NTS and TS represent non-target strand and target strand, respectively.

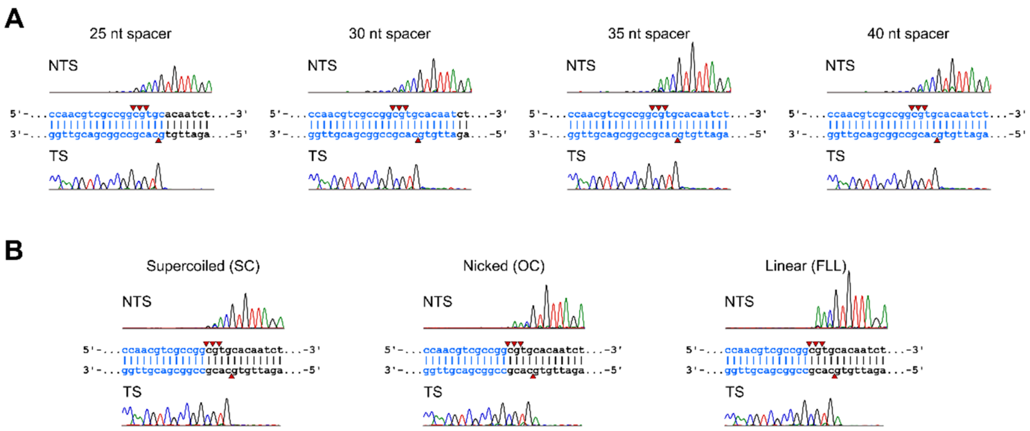


**Supplementary Figure S14**. Run-off sequencing of Cas14a1 cleaved plasmid DNA. Plasmid DNA cleavage resulted in a double-stranded break centered around positions 20-24 bp 3' of the PAM. The cleavage pattern was independent of spacer length (**A**) and plasmid topology (**B**). NTS and TS represent non-target strand and target strand, respectively.

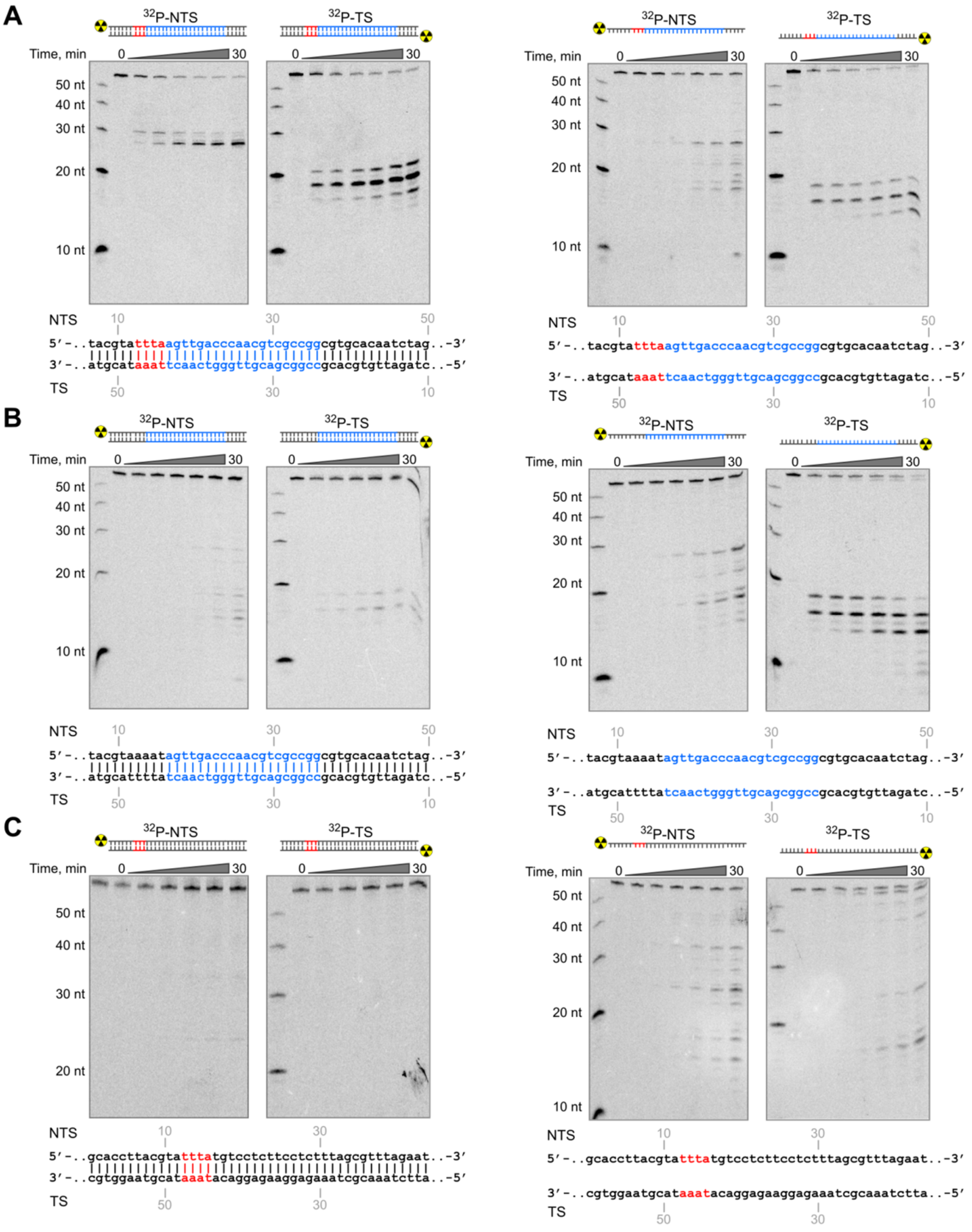


**Supplementary Figure S15**. Oligoduplex cleavage by Cas14a1 RNP complex. (**A**) Purified Cas14a1 RNP complexes cleaved radiolabeled dsDNA oligoduplexes containing a sgRNA target in a PAM-dependent manner generating a staggered cleavage pattern. dsDNA substrates without PAM (**B**) or target sequence (**C**) were not cleaved by the Cas14a1 RNP complex. ssDNA substrates complementary to the sgRNA spacer sequence were also cleaved albeit in the PAM-independent manner (*A* and *B*). NTS and TS represent non-target strand and target strand, respectively.

**Supplementary Data S1 (separate file).** DNA, RNA and protein sequences used in this study.
